# Decreased scene-selective activity within the posterior intraparietal cortex in amblyopic adults

**DOI:** 10.1101/2024.06.05.597579

**Authors:** Sarala N. Malladi, Jan Skerswetat, Roger B.H. Tootell, Eric D. Gaier, Peter Bex, David G. Hunter, Shahin Nasr

## Abstract

Amblyopia is a developmental disorder associated with reduced performance in visually guided tasks, including binocular navigation within natural environments. To help understand the underlying neurological disorder, we used fMRI to test the impact of amblyopia on the functional organization of scene-selective cortical areas, including the posterior intraparietal gyrus scene-selective (PIGS) area, a recently discovered region that responds selectively to ego-motion within naturalistic environments (Kennedy et al., 2024).

Nineteen amblyopic adults (10 female) and thirty age-matched controls (12 female) participated in this study. Amblyopic participants spanned a wide range of amblyopia severity, based on their interocular visual acuity difference and stereoacuity. The visual function questionnaire (VFQ-39) was used to assess the participants’ perception of their visual capabilities.

Compared to controls, we found weaker scene-selective activity within the PIGS area in amblyopic individuals. By contrast, the level of scene-selective activity across the occipital place area (OPA), parahippocampal place area (PPA), and retrosplenial cortex (RSC)) remained comparable between amblyopic and control participants. The subjects’ scores on “general vision” (VFQ-39 subscale) correlated with the level of scene-selective activity in PIGS.

These results provide novel and direct evidence for amblyopia-related changes in scene-processing networks, thus enabling future studies to potentially link these changes across the spectrum of documented disabilities in amblyopia.

## 1. Introduction

Amblyopia is a developmental disorder caused by disruption of balanced binocular input during early life stages. Amblyopic individuals show reduced visual acuity, typically in one eye, despite normal ocular structure. Originally, it was suggested that amblyopic individuals should show performance comparable to that of controls during binocular tasks by relying on the fellow eye. However, emerging evidence suggests that amblyopic children and adults show poorer performance during visually guided activities, even when these tasks were conducted binocularly (Birch, 2013; Kelly et al., 2015; Birch et al., 2019; Birch et al., 2022). Among these impairments, limitations in distance vision and peripheral vision especially affect the self-efficacy and quality of life (QoL) for amblyopic individuals.

According to QoL studies, amblyopic individuals have difficulty participating in outdoor physical activities, navigating around objects without collision, and even crossing streets (Kumaran et al., 2019; Randhawa et al., 2023). Consistent with these reports, amblyopia is also associated with impaired egocentric distance perception either in near personal space (<2 m) (Melmoth and Grant, 2006; Carlton and Kaltenthaler, 2011; Grant and Moseley, 2011) or farther action space (Ooi and He, 2015), even when experiments are conducted binocularly. Moreover, amblyopic subjects show poorer scene discrimination performance, a task the relies on ego-distance encoding, even when images are perceived binocularly (Mirabella et al., 2011).

It could be argued that perceptual impairments in amblyopia arise from poorer depth perception in amblyopic vs. non-amblyopic individuals (McKee et al., 1990; McKee et al., 2003; Levi et al., 2015). According to animal models, impairments in depth perception are associated with a decrease in the number of binocularly responsive neurons in V1 (Crawford and Von Noorden, 1979; Horton et al., 1997; Smith III et al., 1997a) and decreased sensitivity to binocular disparity within cortical areas V1-V3A (Kumagami et al., 2000; Bi et al., 2011). Although these neuronal deficits are expected to impact the functional development of higher-level (“downstream”) visual areas that are involved in complex cognitive tasks such as scene perception and/or ego-motion control, there is no direct evidence to support this hypothesis, to the best of our knowledge.

In humans, there is a network of visual areas that shows a selectively higher response to scenes, when compared to other visual object categories (Nasr et al., 2011; Kennedy et al., 2024). In this study, we test the hypothesis that amblyopia preferentially impacts function in one or more of these scene-selective area(s). These areas include (but are not limited to): (*i*) the temporal place area known as the parahippocampal place area (PPA) (Epstein and Kanwisher, 1998), (*ii*) the occipital place area (OPA)(Grill-Spector, 2003; Dilks et al., 2013), (*iii*) the medial place area located near the retrosplenial cortex (RSC) (Maguire, 2001; Park and Chun, 2009), and (*iv*) the posterior intraparietal gyrus scene-selective area (PIGS). Compared to other category-selective visual areas, activity within the scene-selective areas relies heavily on the distance between the visual objects and the observer (Kravitz et al., 2011; Persichetti and Dilks, 2016; Park and Park, 2020). Among these areas, PIGS (Kennedy et al., 2024) and OPA (Kamps et al., 2016; Jones et al., 2023) also respond selectively to ego-motion within naturalistic scenes. Thus, considering impairments in distance and ego-motion perception among amblyopic individuals, we expected the amblyopia impact to be stronger on the scene-selective areas, especially in PIGS and OPA.

To test the hypothesis that amblyopia influences the function of scene-selective areas, we first tested (and confirmed) previous reports that amblyopic individuals self-report lower scores for general vision, distance activities, and peripheral vision, compared to age-matched controls (Kumaran et al., 2019; Randhawa et al., 2023). Next, we used fMRI to compare the evoked scene- and object-selective activity between amblyopic individuals and controls. To reduce the impact of decreased stereoacuity (a common impairment among amblyopic individuals) on the evoked brain response, participants were presented with binocular, 2D images during the fMRI scabs. Results of these scans showed a decreased scene-selective activity in area PIGS (but not the other scene-selective areas) in amblyopic individuals compared to controls.

## 2. Methods

### 2.1. Participants

Forty-nine adult humans, aged 18-54 years, participated in this study. Among them, nineteen individuals (10 females) were diagnosed with amblyopia. The remaining thirty participants (15 females) had normal corrected visual acuity in both eyes. All participants had radiologically normal brains, without any history of neuropsychological disorder. All experimental procedures conformed to NIH guidelines and were approved by Massachusetts General Hospital protocols. Written informed consent was obtained from all subjects before the experiments.

### 2.2. General procedure

The study consisted of a behavioral experiment and two neuroimaging tests. The behavioral tests were performed outside of the scanner, including 1) answering a questionnaire, and 2) conducting ophthalmological assessments. The neuroimaging experiments were conducted on a different day relative to the behavioral tests, inside of a 3T scanner. As demonstrated in **Tables Table *1***, a subset of volunteers participated in all experiments. The others participated in 1 or 2 experiments, depending on their availability at the time.

**Table 1.** Participants demography and contributions.

| ID | Age | Gende | Exp 1 | Exp 2 | Exp 3 |  | ID | Age | Sex | Exp 1 | Exp 2 | Exp 3 |
| --- | --- | --- | --- | --- | --- | --- | --- | --- | --- | --- | --- | --- |
| Controls |  |  |  |  |  |  | Controls |  |  |  |  |  |
| C1 | 24 | M | 1 | 1 | 1 |  | C16 | 31 | F | 0 | 1 | 1 |
| C2 | 30 | M | 1 | 1 | 1 |  | C17 | 28 | M | 0 | 1 | 1 |
| C3 | 26 | F | 1 | 1 | 0 |  | C18 | 30 | M | 0 | 1 | 1 |
| C4 | 32 | M | 1 | 1 | 0 |  | C19 | 26 | F | 0 | 1 | 1 |
| C5 | 22 | F | 1 | 1 | 0 |  | C20 | 22 | M | 0 | 1 | 1 |
| C6 | 43 | M | 1 | 0 | 0 |  | C21 | 25 | F | 0 | 1 | 1 |
| C7 | 38 | F | 1 | 0 | 0 |  | C22 | 28 | F | 0 | 1 | 1 |
| C8 | 38 | F | 1 | 0 | 0 |  | C23 | 30 | M | 0 | 1 | 1 |
| C9 | 35 | M | 1 | 0 | 0 |  | C24 | 32 | F | 0 | 1 | 1 |
| C10 | 26 | M | 1 | 0 | 0 |  | C25 | 30 | F | 0 | 1 | 1 |
| C11 | 23 | F | 1 | 0 | 0 |  | C26 | 25 | F | 0 | 1 | 1 |
| C12 | 34 | F | 0 | 1 | 1 |  | C27 | 40 | F | 0 | 1 | 0 |
| C13 | 32 | F | 0 | 1 | 1 |  | C28 | 26 | M | 0 | 1 | 0 |
| C14 | 38 | M | 0 | 1 | 1 |  | C29 | 29 | M | 0 | 1 | 0 |
| C15 | 37 | M | 0 | 1 | 1 |  | C30 | 27 | M | 0 | 1 | 0 |
| Strabismic |  |  |  |  |  |  | Anisotropic / Deprivational |  |  |  |  |  |
| S1 | 40 | F | 1 | 1 | 1 |  | A1 | 31 | F | 1 | 1 | 1 |
| S2 | 20 | M | 1 | 1 | 1 |  | A2 | 23 | M | 1 | 1 | 1 |
| S3 | 28 | M | 1 | 1 | 1 |  | A3 | 35 | M | 1 | 1 | 1 |
| S4 | 26 | F | 1 | 1 | 1 |  | A4 | 20 | F | 1 | 1 | 1 |
| S5 | 31 | F | 1 | 1 | 1 |  | A5 | 26 | M | 1 | 1 | 1 |
| C6 | 26 | F | 1 | 1 | 1 |  | A6 | 19 | F | 1 | 1 | 1 |
| S7 | 21 | F | 1 | 1 | 1 |  | A7 | 24 | F | 1 | 1 | 1 |
| S8 | 56 | M | 1 | 0 | 0 |  | A8 | 26 | M | 0 | 1 | 1 |
| S9 | 28 | M | 0 | 1 | 1 |  | A9 | 18 | F | 0 | 1 | 1 |
|  |  |  |  |  |  |  | D1 | 30 | M | 1 | 1 | 1 |

#### 2.2.1. Experiment 1 – behavioral tests

Among the participants, sixteen amblyopic individuals (9 females) and eleven controls (5 females) participated in the behavioral experiment (**Tables Table *1***). This experiment consisted of two parts: The first part was based on the National Eye Institute (NEI) visual function questionnaire (NEI-VFQ 39), which is a well-validated questionnaire on visual function and disabilities, including subscales on general vision, ocular pain, near vision, distance vision, vision-specific social function, vision-specific mental health, vision-specific role function, dependency, driving, peripheral vision, and color vision.

The second set of behavioral measurements included ophthalmological tests that were conducted outside of the scanner by an optometrist (JS) with extensive experience evaluating amblyopic individuals. Those measurements assessed each participant’s best-corrected monocular distance visual acuity, binocular visual acuity (ETDRS retro luminant chart (Precision Vision)), the presence of peripheral monocular suppression (Worth 4-dot at near), and stereoacuity (Randot stereo test (Stereo Optical).

#### 2.2.2. Experiment 2 – scene-selective activity measurement

Eighteen amblyopic individuals, plus twenty-four controls, participated in this fMRI experiment. During the MRI scans, participants were presented with 8 naturally colored images of real-world scenes vs. group faces (Nasr et al., 2011; Kennedy et al., 2024). Scene and face stimuli were retinotopically centered and subtended 20° × 26° of visual field, without any significant differences between their root mean square (RMS) contrast (t(14) =1.10, *p*=0.29). Importantly, we recently showed that: (a) PIGS responds selectively to a wide range of scenes (including indoor and outdoor scene) without any apparent change in the center of activity, and (b) the PIG responses remained comparable to ‘scene vs. face’ and ‘scene vs. object/face’ contrasts (Kennedy et al., 2024).

Scene and face stimuli were presented in different blocks (16 s per block and 1 s per image). Each subject participated in 6 runs. Each run consisted of 10 blocks, plus 32 s of a blank gray presentation at the beginning and at the end of each block. Within each run, the sequence of blocks and images within each block were randomized. Data from one amblyopic subject was excluded due to a technical problem in stimulus presentation.

During the scan, stimuli were presented via a projector (1024 × 768 pixel resolution, 60 Hz refresh rate) onto a rear-projection screen. Subjects viewed the stimuli through a mirror mounted on the receive coil array. To ensure that subjects were attending to the screen, each subject was instructed to report color changes (red to blue and vice versa) of a centrally presented fixation point (0.1° × 0.1°) by pressing a key on the keypad. Subject detection accuracy remained above 75% and showed no significant difference in color change detection performance across experimental conditions (*p*>0.10). MATLAB (MathWorks; Natick, MA, USA) and the Psychophysics Toolbox (Brainard, 1997; Pelli, 1997) were used to control stimulus presentation.

#### 2.2.3. Experiment 3 – object-selective activity measurement

Among those who participated in Experiment 2, eighteen amblyopic individuals and seventeen controls (9 females) agreed to be scanned further to measure their response to intact and scrambled objects. Stimuli consisted of 38 gray-scale images of intact everyday objects (e.g., tools, furniture and fruits) and their scrambled versions (i.e. no RMS contrast difference) (Nasr et al., 2013; Yue et al., 2013). Stimuli were retinotopically centered on a fixation spot and subtended 20° × 20° of visual field. Intact and scrambled images were presented in different blocks (16 s per block and 1 s per image). Each subject participated in 6 runs, and each run consisted of 8 blocks plus 16 s of blank gray presentation at the beginning and at the end of each block. Within each run, the sequences of blocks (and images within blocks) were randomized. Other details of the stimulus presentation and the participant’s task during the experiments were identical to Experiment 2.

### 2.3. Imaging

Subjects were scanned in a horizontal 3T scanner (Tim Trio, Siemens Healthcare, Erlangen, Germany). Gradient echo EPI sequences were used for functional imaging. Functional data were acquired using single-shot gradient echo EPI, using isotropic voxels, nominally 3.0 mm on each side (TR=2000 ms; TE=30 ms; flip angle=90°; band width (BW)=2298 Hz/pix; echo-spacing= 0.5 ms; no partial Fourier; 33 axial slices covering the entire brain; and no acceleration). During the scan session, structural (anatomical) data were also acquired for each subject using a 3D T1-weighted MPRAGE sequence (TR=2530 ms; TE=3.39 ms; TI=1100 ms; flip angle=7°; BW=200 Hz/pix; echo-spacing=8.2 ms; voxel size=1.0×1.0×1.33 mm).

### 2.4. Data Analysis

Structural and functional data analysis were conducted based on using FreeSurfer (Fischl, 2012).

#### 2.4.1. Structural data analysis

For each subject, inflated and flattened cortical surfaces were reconstructed based on the high-resolution anatomical data (Dale et al., 1999; Fischl et al., 1999; Fischl et al., 2002). A pial ‘surface’ was generated relative to the gray matter boundary with the surrounding cerebrospinal fluid or CSF (i.e., the GM-CSF interface). A white matter surface was also generated as the interface between white and gray matter (i.e. WM-GM interface). An additional (“midlayer”) surface was also generated at 50% of the depth of the local gray matter (Dale et al., 1999).

#### 2.4.2. Individual-level functional data analysis

All functional data were rigidly aligned (6 df) relative to subject’s own structural scan, using rigid Boundary-Based Registration (Greve and Fischl, 2009), followed by motion correction. That data was spatially smoothed using a 3D Gaussian kernel (2 mm FWHM). Subsequently, a standard hemodynamic model based on a gamma function was fitted to the fMRI signal, sampled from the middle of cortical gray matter (defined for each subject based on their structural scan), to estimate the amplitude of the BOLD response. Finally, vertex-wise statistical tests were conducted by computing contrasts based on a univariate general linear model (Friston et al., 1999). For presentation of activity maps based on individual subjects, the resultant significance maps were projected onto a common human brain template (fsaverage; (Fischl, 2012).

#### 2.4.3. Group-level functional data analysis

To generate group-averaged maps, functional maps were spatially normalized across subjects, then averaged using random-effects models and corrected for multiple comparisons (Friston et al., 1999). The resultant significance maps were projected onto the fsaverage.

#### 2.4.4. Vertex-wise between-groups comparison

Unless otherwise indicated, between-group (amblyopic vs. control participants) activity difference maps were also generated based on a random-effects model, after correcting for multiple comparisons.

#### 2.4.5. Region of interest (ROI) analysis

The main ROIs included area PIGS, plus each of the three previously known scene-selective areas (PPA, RSC, OPA). Each of these ROIs was defined independently for amblyopic and control individuals, based on the corresponding group-averaged activity maps. Specifically, for each group, the group-averaged ‘scene > face’ activity map (based on random-effects and after correction for multiple comparisons) was generated independently based on the odd and even runs. For amblyopic and control participants, the ROIs were localized at *p<*10^-2^ and *p*<0.05 threshold levels (respectively). These ROIs were then projected back into each individual’s native space.

The ROIs that were defined based on the odd runs were used to measure the level of activity evoked during the even runs, and vice versa. This procedure assured us that the ROIs were defined based on an independent dataset compared to the test data.

In addition to the scene-selective areas, we also used the lateral object-selective complex (LOC; (Grill-Spector et al., 2001)) and the area V6 (Pitzalis et al., 2010) as control ROIs. Area LOC was localized functionally (Experiment 3), based on the group-averaged activity map evoked in response to “intact > scrambled objects” functional contrast. Here again, ROIs were defined independently for each group, based on the corresponding group-averaged activity maps. Area V6 was localized using a probabilistic label generated previously based on an independent group of individuals (Kennedy et al., 2024).

#### 2.4.6. Comparing the size of scene-selective areas

To compare the size of scene-selective areas between amblyopic individuals and controls, these areas were localized for each subject based on their own scene-selective activity map at a threshold of *p*<10^-2^. These measurements were then normalized relative to the size of the entire cerebral cortex. This procedure assured us that our tests were not confounded by differences in overall brain size.

### 2.5. Statistical tests

To test the effect of independent parameters, we applied paired t-tests and/or a repeated-measures ANOVA, with Greenhouse-Geisser correction whenever the sphericity assumption was violated. The effect of group was tested by comparing the response from controls vs. amblyopic individuals, irrespective of the amblyopia sub-type, unless otherwise is noted. All results were corrected for multiple comparisons.

### 2.6. Data sharing statement

All data, codes and stimuli are ready to be shared upon request. MATLAB (RRID: SCR_001622; https://www.mathworks.com).

FreeSurfer (RRID:SCR_001847; https://surfer.nmr.mgh.harvard.edu/fswiki/FsFast).

Psychophysics Toolbox (RRID:SCR_002881; http://psychtoolbox.org/docs/Psychtoolbox).

## 3. Results

### 3.1. Participants age and gender distribution

Nineteen amblyopic individuals (10 females), aged 18-56 years, and thirty controls (15 females) with normal or corrected-to-normal visual acuity, aged 22-43 years, participated in this study (**Tables Table *1***). Amblyopic participants included 9 with strabismus, 9 with anisometropia, and 1 with deprivational amblyopia. No participants had combined strabismic and anisometropic amblyopia. Independent applications of t-tests did not yield any significant age differences between amblyopic vs. control participants (t(48)=1.18, *p*=0.25) or between anisometropic vs. strabismic participants (t(17)=1.38, *p*=0.19). Application of this analysis to the subset of subjects who participated in each experiment yielded the same result. Thus, potential differences between groups could not be attributed solely to age differences.

### 3.2. Experiment 1 – Ophthalmologic and VFQ-39 tests

Sixteen amblyopic individuals and 11 controls were examined by an optometrist with extensive experience with amblyopia to measure their monocular and binocular visual acuity, monocular suppression, and stereoacuity. They also answered the VFQ-39 questionnaire (see Methods and **Table *1***).

#### 3.2.1. Ophthalmological assessment

All (except for one) amblyopic individuals showed evidence for either monocular suppression or diplopia (Worth 4-Dot; **Table *2***). In contrast, all tested controls showed binocular fusion. Moreover, compared to controls, amblyopic individuals showed a significantly higher interocular visual acuity difference (t(25)=3.15; *p*<0.01) and poorer stereoacuity (t(25)=2.52; *p*<0.01), as expected. However, binocular visual acuity did not differ significantly between the two groups (t(25)=1.27; *p*=0.22). Amblyopia severity (assessed by the interocular visual acuity difference) was comparable between the anisomteropic (n=7) and strabismic (n=8) individuals (t(13)=1.61; *p*=0.13). Consistent with previous reports (McKee et al., 2003; Levi et al., 2015), more strabismic individuals demonstrated severely impaired stereoacuity (>800 arc seconds) than anisometropic individuals. However, stereoacuity was statistically comparable between anisometropic and strabismic participants (t(13)=1.57; *p*=0.14).

**Table 2.** Ophthalmologic assessment of all participants.

| ID | Age of Diagnosis | Right Eye Visual Acuity | Left Eye Visual Acuity | Binocular Visual Acuity | Fellow/Dominant Eye | Suppression Worth 4Dots | Randot Stereoacuity | Angle of Strabismus (at 4 m in PD) |
| --- | --- | --- | --- | --- | --- | --- | --- | --- |
| S1 | <1 | -0.06 | +0.06 | -0.06 | RE | Diplopia | >500 | 16/25 |
| S2 | 3 | +0.09 | +0.30 | +0.09 | LE | RE | >500 | 12/10 |
| S3 | 6 | +0.48 | -0.02 | +0.02 | LE | RE | >500 | 25/18 |
| S4 | 6 | +0.00 | -0.06 | +0.00 | LE | Diplopia | 50 | 4/4 |
| S5 | 3 | +0.26 | +0.04 | +0.04 | LE | RE | >500 | 10/8 |
| S6 | 2 | +0.46 | -0.06 | -0.14 | LE | RE | >500 | 10/8 |
| S7 | 4 | -0.08 | -0.10 | -0.08 | LE | Diplopia | 70 | 20/20 |
| S8 | 5 | -0.20 | +0.06 | +0.06 | RE | LE | >500 | 16/16 |
| S9 | 2 | +0.00 | +0.60 | +0.00 | RE | LE | >500 | 14/10 |
| A1 | 5 | -0.22 | +0.20 | -0.16 | RE | None | >500 | None |
| A2 | 5 | -0.08 | +0.30 | -0.04 | RE | None | 400 | None |
| A3 | 11 | -0.04 | +0.26 | -0.04 | RE | LE | 200 | None |
| A4 | 6 | +0.06 | +0.32 | +0.00 | RE | None | 40 | None |
| A5 | 8 | +0.00 | +0.17 | +0.00 | RE | Diplopia | 200 | None |
| A6 | 5 | +1.00 | -0.08 | -0.08 | LE | RE | >500 | None |
| A7 | 8 | +0.64 | -0.10 | -0.10 | LE | RE | >500 | None |
| A8 | 6 | -1.25 | -0.25 | -0.25 | RE | LE | >500 | None |
| A9 | 7 | +0.40 | +0.00 | +0.00 | RE | None | >500 | None |
| D1 | 4 | -0.26 | +0.10 | -0.26 | RE | LE | 100 | None |

#### 3.2.2. Qualitative assessment of amblyopia impacts on visual capabilities

Previous studies reported that amblyopic individuals struggle with distance activities and peripheral vision (Kumaran et al., 2019; Randhawa et al., 2023). To directly test for analogous results in our cohort, participants in Experiment 1 received the VFQ-39 questionnaire. Consistent with previous studies, we found significantly lower (i.e. worse) scores for amblyopic individuals compared to controls in the general vision, distance activities, and peripheral vision categories (*p*≤0.01; **Table 3**). In contrast, we did not find any significant difference between the two groups in near activities, color vision, or driving capabilities *(p≥*0.08). No significant differences were found between the strabismic and anisometropic individuals across the VFQ subscales (*p≥*0.14).

**Table 3.** VFQ-39 subscales.

| **Subscale** |  | **Controls (n=11)**<br>**(mean ± S.D.)** | **Strabismic (n=8)**<br>**(mean ± S.D.)** | **Anisometric (n=8)**<br>**(mean ± S.D.)** | **Control vs.**<br>**Amblyopic**<br>**participants<sup>a</sup>**<br>**(p-value)** |
| --- | --- | --- | --- | --- | --- |
| General Health | | 88.18 ± 12.50 | 75.31 ± 9.20 | 77.81 ± 15.09 | 0.02* |
| General Vision | | 85.45 ± 10.83 | 69.38 ± 16.57 | 66.25 ± 9.16 | <10<sup>-3</sup>** |
| Ocular Pain | | 94.32 ± 10.25 | 90.62 ± 18.60 | 87.50 ± 11.57 | 0.32 |
| Near Activities | | 98.86 ± 3.77 | 97.40 ± 4.42 | 92.71 ± 7.30 | 0.09 |
| Distance Activities | | 97.35 ± 5.36 | 86.98 ± 13.63 | 86.98 ± 8.16 | <0.01** |
| Vision Specific | Social Functioning | 100.00 ± 0.00 | 95.83 ± 8.91 | 98.96 ± 2.95 | 0.21 |
| | Mental Health | 97.27 ± 3.44 | 86.25 ± 19.41 | 83.75 ± 12.17 | 0.02* |
| | Role Difficulties | 100.00 ± 0.00 | 92.97 ± 9.70 | 93.75 ± 4.72 | <0.01** |
| | Dependency | 98.86 ± 3.77 | 97.66 ± 4.65 | 100.00 ± 0.00 | 0.98 |
| Driving | | 84.72 ± 13.66 | 77.08 ± 21.53 | 73.44 ± 14.07 | 0.16 |
| Color Vision | | 100.00 ± 0.00 | 100.00 ± 0.00 | 100.00 ± 0.00 | 1.00 |
| Peripheral Vision | | 97.73 ± 7.54 | 81.25 ± 22.16 | 84.38 ± 12.94 | 0.01* |

**α:** The effect of group was tested between controls vs. amblyopic individuals, irrespective of their amblyopia sub-type. A separate test did not show any significant differences between anisomteropic and strabismic individuals (not shown here).

#### 3.2.2. Predictability of VFQ-39 scores based on the ophthalmological measurements

We tested whether the VFQ-39 scores for general vision, distance activities and peripheral vision were predictable based on the level of interocular visual acuity difference, binocular visual acuity and stereoacuity. Among these VFQ-39 subscales, separate Pearson correlation tests showed a significant linear relationship between the interocular visual acuity difference and general vision (R^2^=0.28; *p*<0.01), and peripheral vision (R^2^=0.17; *p*=0.03). We also found a marginal (statistically non-significant) correlation between the binocular visual acuity and general vision (R^2^=0.13; *p*=0.06). The correlations between the other factors were non-significant (R^2^<0.10; *p*>0.11).

### 3.3. Experiment 2 – Scene-selective cortical response

Experiment 2 was designed to test whether amblyopia is associated with a decrease in the amplitude of scene-selective responses. Eighteen amblyopic individuals, plus twenty-four controls, participated in this experiment (**Table *1***) and were presented with scene and face stimuli in different blocks (see Methods).

#### 3.3.1. Head position stability

Head motion has a strong impact on the fMRI signal, and it may influence the level and pattern of evoked fMRI responses, which might thus confound between-group comparisons. However, a t-test applied to the measured level of head motion did not yield a significant difference between the two groups (t(40)=1.58, *p*=0.12). Thus, head motion was statistically comparable across the two groups. Nevertheless, head motion was included as a nuisance co-variate in all analyses, to reduce any residual impact of head motion on our findings.

#### 3.3.2. Group-level localization of scene-selective areas

Figure 1A-B shows the scene-selective activity maps generated by contrasting the evoked response to scene vs. face stimuli in control and amblyopic subjects, respectively. In both groups, we were able to identify the PPA, RSC and TPS/OPA, without any apparent differences in the location of these areas between control and amblyopic groups. In controls, we were also able to detect area PIGS, close to the posterior border of the parieto-occipital sulcus within the intraparietal gyrus (Figure 1A and Figure 2A) (Kennedy et al., 2024). In the group-averaged maps from the amblyopic subjects, PIGS was detectable only when the threshold was lowered to *p*<0.05 (Figure 2B). Even in these low-threshold maps, the center of PIGS appeared to be located more ventrally in amblyopic participants compared to the control group. This difference was also detectable when we generated the PIGS label based on the group-averaged activity maps evoked independently during odd and even runs (Figure 3 **– Consistency of localization in scene-selective areas across odd and even runs. In amblyopic and control groups, scene-selective areas were detected across odd and even runs** Figure 3).

**Figure 1.**
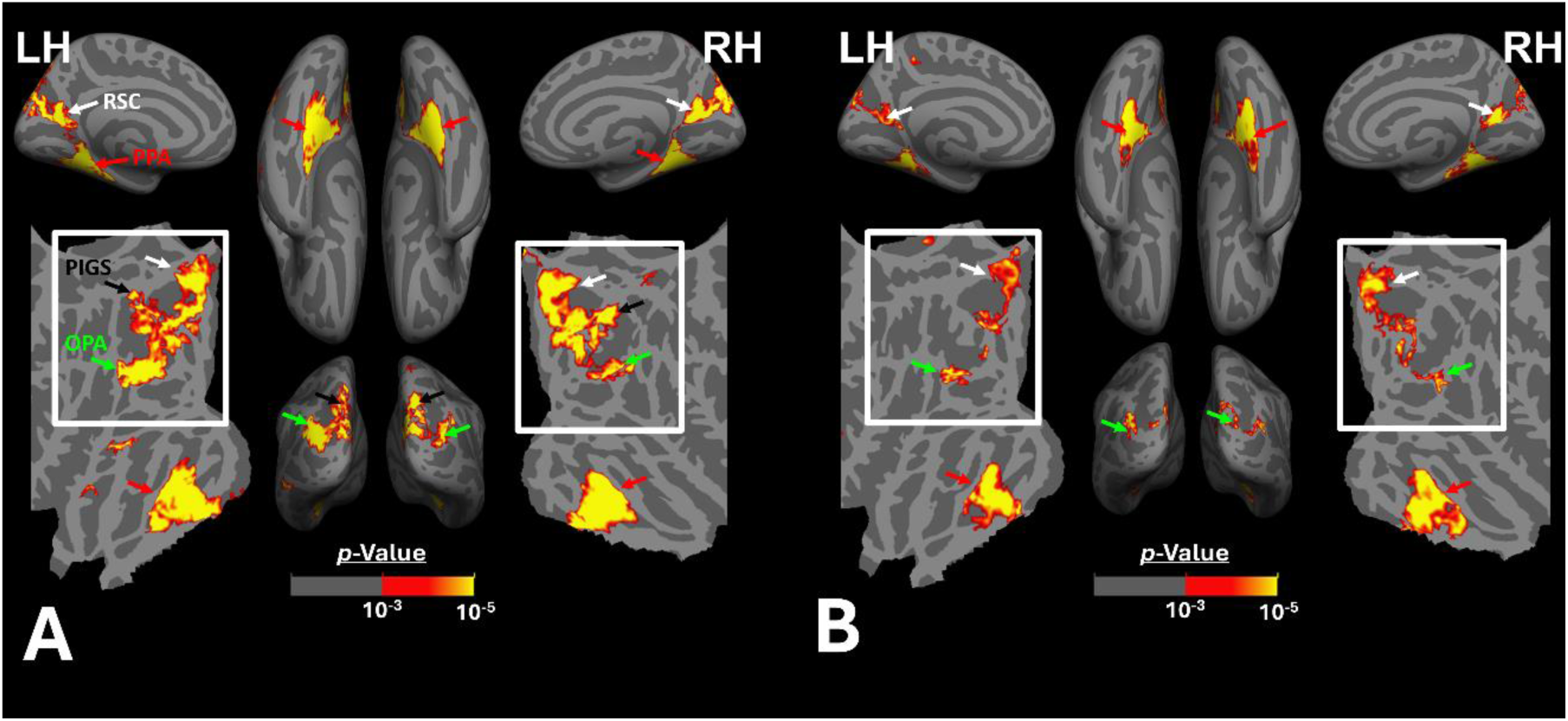
Group-averaged scene-selective activity in control (A) and amblyopic (B) participants. By measuring the response to “scene > face” contrast, we located areas PPA, RSC, OPA and PIG (indicated by red, white, green and black arrowheads respectively) for controls. In amblyopic individuals, we detected the same overall scene-selective activity pattern. However, in amblyopic compared to control participants, we found a weaker scene-selective activity within the posterior intraparietal gyrus (see also Figure 2). Both group-averaged activity maps were calculated based on random-effect analyses, and were overlaid on a common brain template (fsaverage). The white inset indicates the occipito-parietal region in which PIGS, OPA and RSC are located.

**Figure 2.**
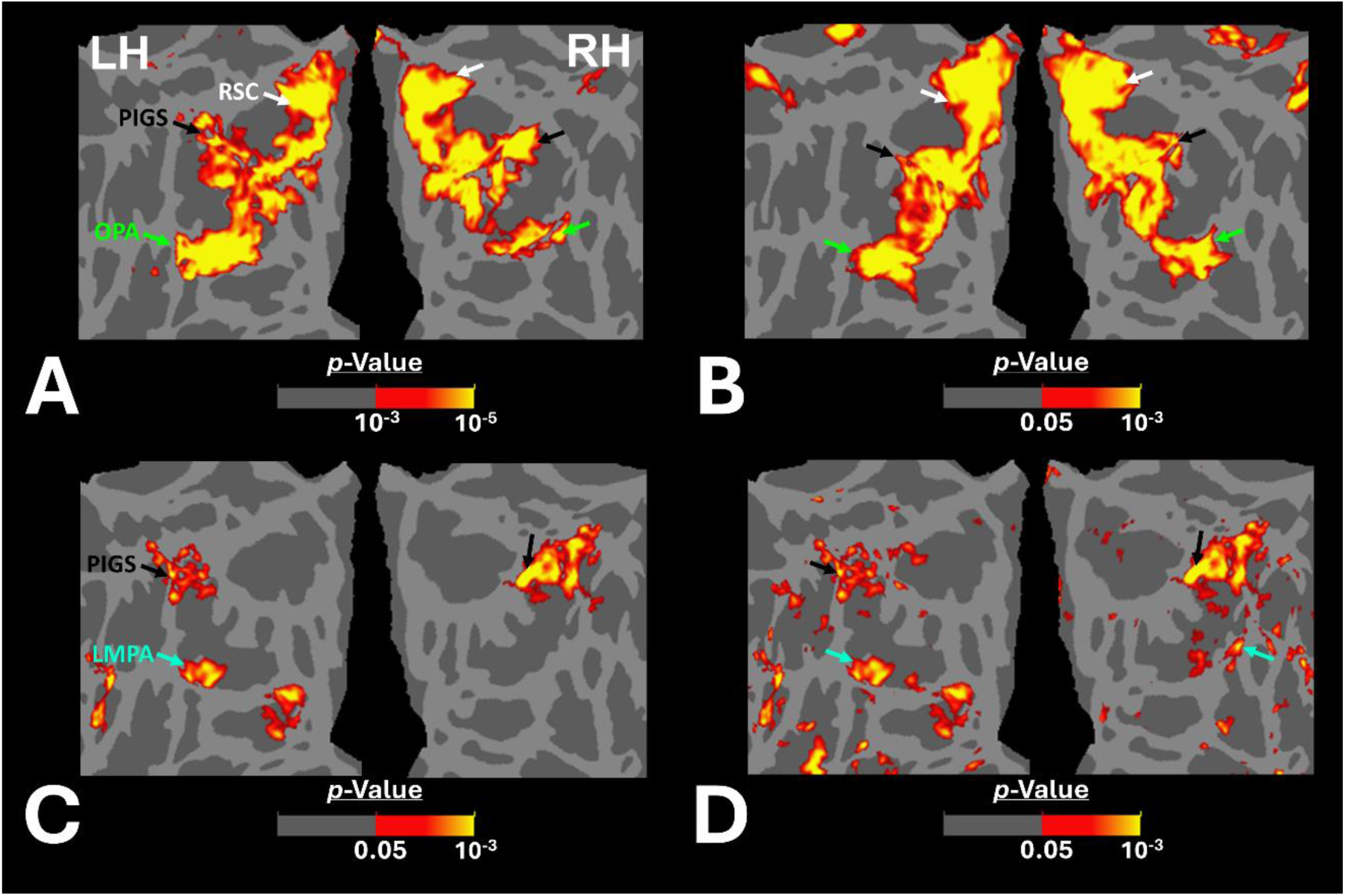
Group-averaged scene-selective activity in amblyopic and control participants across the occipito-parietal region. Panels A and B show the activity maps in controls and amblyopic individuals, respectively. For amblyopic individuals, the activity map is generated based on lower threshold levels. Despite using those more liberal thresholds, the scene-selective activity within the posterior intraparietal gyrus appeared to be weaker in amblyopic compared to control participants. Panels C and D show the between-group scene-selective activity differences, with and without correction for multiple comparisons, respectively. Consistently, we found bilateral scene-selective activity difference within the posterior intraparietal region. In Panel D, beyond the sensory scene-selective areas, we also noticed a bilateral activity difference within the LPMA region, as reported previously using conventional fMRI (Steel et al., 2021; Steel et al., 2023). Other details are similar to those in Figure 1.

**Figure 3.**
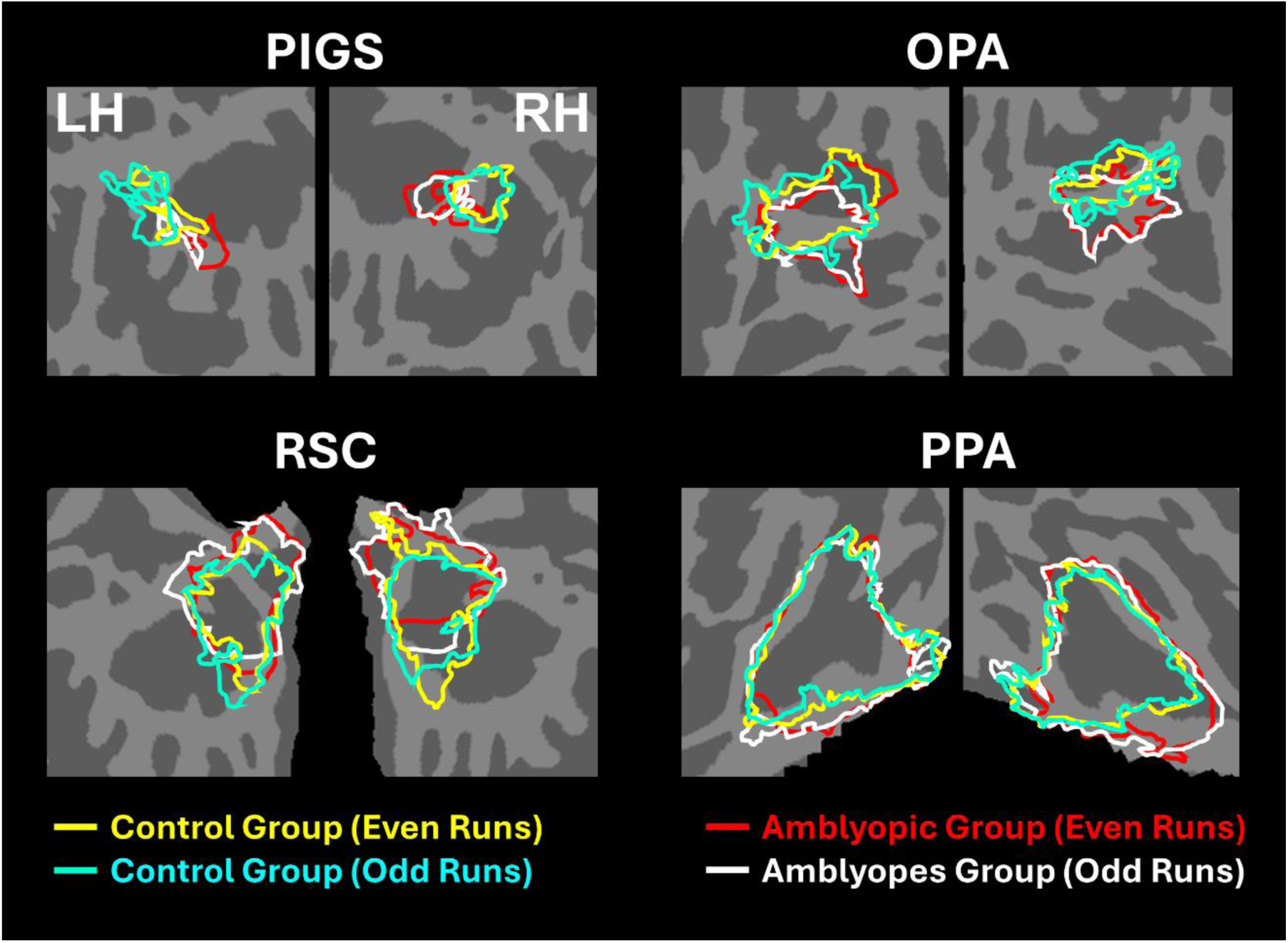
Consistency of localization in scene-selective areas across odd and even runs. In amblyopic and control groups, scene-selective areas were detected across odd and even runs. PIGS in amblyopic individuals is on average located more ventrally compared to controls, whereas RSC, OPA and PPA are located in a similar location between amblyopic and control participants. This difference in PIGS location was detected during odd and even runs, without any apparent differences.

Figure 2C shows the vertex-wise map of activity difference between controls and amblyopic individuals, after correction for multiple comparisons. In both hemispheres, we found stronger scene-selective activity within PIGS in the controls compared to amblyopic participants. Besides PIGS, we also found a significant between-group activity difference in the anterior intraparietal gyrus, on the opposite side relative to PIGS, and posteriorly relative to the medial and superior temporal sulci. Named lateral place memory area (LPMA), this region is expected to be involved in place memory retrieval (Steel et al., 2021; Steel et al., 2023). This latter activity appeared to be stronger in the left compared to the right hemisphere. However, the same pattern of activity was also detectable in the right LPMA region, when analyzed without correction for multiple comparisons (Figure 2D).

Beyond the sensory areas, we also found bilateral activity differences within the temporal parietal junction (TPJ) and dorsolateral prefrontal cortex (DPFC), two regions that are expected to be involved in attention control and decision making (Figure 4). Importantly, these activity differences were detected even though subjects were not instructed to do any scene-related tasks such as memory recall (Steel et al., 2021; Steel et al., 2023) or spatial comparison (Nasr and Tootell, 2012; Nasr et al., 2013). The same areas also did not appear within the scene-selectivity maps in either control or amblyopic groups (Figure 1A-B). Thus, these differential activities were most likely due to subthreshold ongoing processes during the blocks. Due to such uncertainties in the nature of these processes, they were excluded from the rest of the data analysis (but see Discussion).

**Figure 4.**
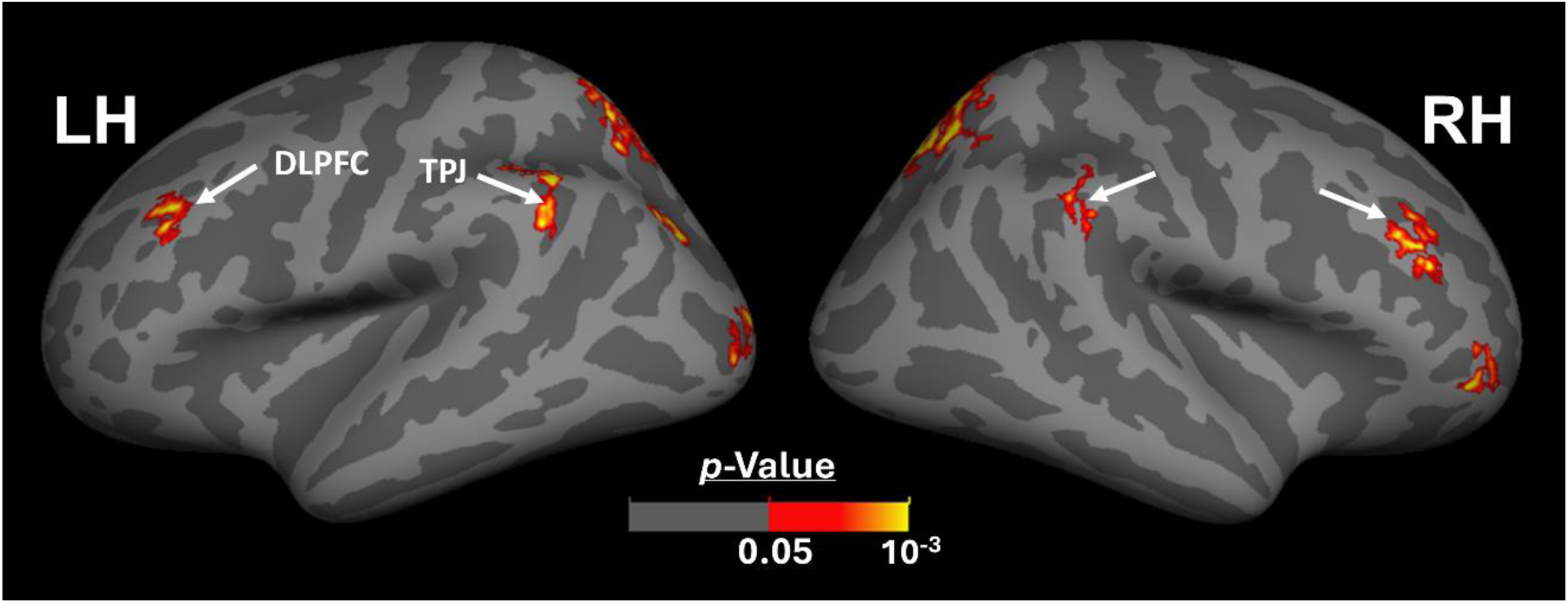
Between-group activity differences outside the visual areas. Beyond the visual areas, we found bilateral scene-selective activity differences between control and amblyopic individuals in areas TPJ and DLPFC. This result suggests that the impact of amblyopia on the response to the ‘scene>face’ contrast may extend to association brain areas. The other details are similar to (Figure 1).

#### 3.3.3. Localization of scene-selective areas in individual subjects

In all participants, including the amblyopic individuals, we were able to localize PPA, RSC, and OPA bilaterally, at a threshold level of *p*<0.01 (Figure 5). PIGS was also detected in all tested control subjects, bilaterally. However, in amblyopic participants, in 8 out of 36 hemispheres, PIGS was *not* detectable at this threshold level. When normalized relative to the size of the whole cortex, the average size of PIGS (but not the other scene-selective areas) was significantly smaller in amblyopic individuals compared to controls (**Table 4**).

**Figure 5.**
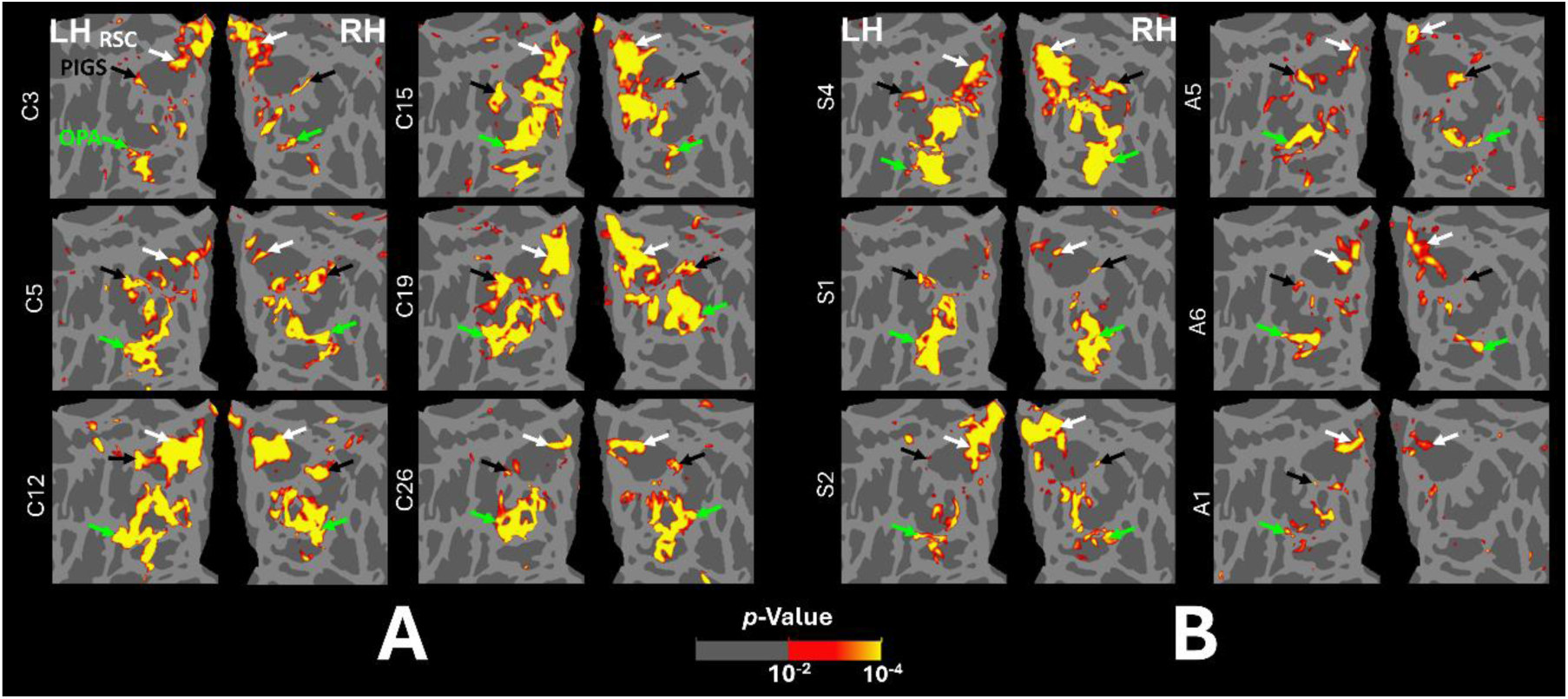
Localization of scene-selective areas within the occipito-parietal cortex across controls and amblyopic individuals. Panel A shows the data from left and right hemispheres of 6 control participants. Panel B shows the data 3 exemplar strabismic (left) and 3 exemplar anisometropic (right) individuals that show strong (top), medium (middle) and weak (bottom) scene-selective activity within the posterior intraparietal region (black arrow). For all subjects, the activity maps were overlaid on a common brain template (fsaverage) to facilitate the comparison across individuals. PIGS, RSC, and OPA are indicated with black, white and green arrowheads.

**Table 4.** Normalized^α^ size of scene-selective areas in amblyopic and control participants.

| Area | Amblyopic<br>LH | Amblyopic<br>RH | Control<br>LH | Control<br>RH | F-value | p-value<br>with<br>correction |
| --- | --- | --- | --- | --- | --- | --- |
| **PIGS** | 0.14 %± 0.14% | 0.14% ± 0.13% | 0.28% ± 0.14% | 0.23% ± 0.12% | 10.12 | 0.01 |
| **OPA** | 0.87% ± 0.51% | 0.75% ± 0.48% | 1.13% ± 0.48% | 0.93% ± 0.51% | 0.63 | >0.99 |
| **RSC** | 0.35% ± 0.36% | 0.40% ± 0.35% | 0.44% ± 0.27% | 0.45% ± 0.45% | 2.34 | 0.52 |
| **PPA** | 1.33% ± 0.46% | 1.17% ± 0.42% | 1.41% ± 0.43% | 1.30% ± 0.49% | 0.65 | >0.99 |

**α:** All values are measured in percentage relative to the overall size of the cortex

#### 3.3.4. The amplitude of the scene-selective activity across amblyopic and control subjects

Figure 6 shows the amplitude of the scene-selective (scene – face) response, evoked within PIGS, OPA, RSC, and PPA. Importantly, the targeted ROIs were determined for amblyopic and control participants independently based on their own group-averaged activity map (see Methods). Also, to avoid logical circularity, the group-averaged activity maps based on the “odd” runs (Figure 3) were used to localize the ROIs that were used to measure the response during “even” runs, and vice versa.

**Figure 6.**
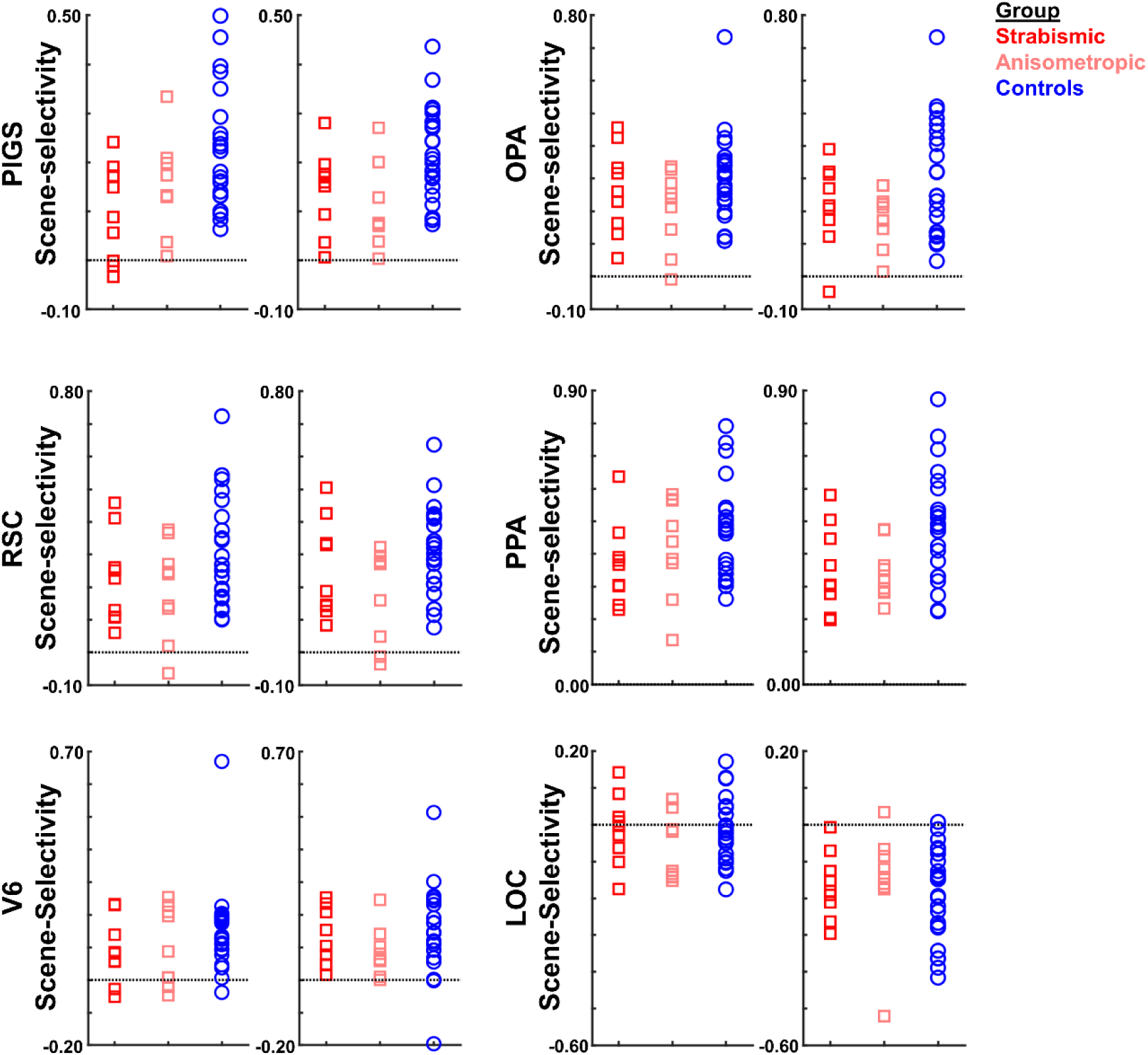
The level of scene-selective activity (Scenes – Faces) was measured across areas PIGS, OPA, RSC, PPA, V6 and LOC. Across all ROIs, we found a significant difference only in the level of scene-selective activity between amblyopic individuals vs. controls in area PIGS. The difference between strabismic and anisometropic individuals remained non-significant across all tested ROIs. For each area, the left and right panels show the activity measured within the left and right hemispheres, respectively. In each panel, each point represents data from one individual participant.

Consistent with the group-averaged activity maps, one-way repeated measures ANOVAs showed a significantly weaker scene-selective activity in PIGS for amblyopic individuals (irrespective of amblyopia subtype) compared to controls (F(40, 1)=12.38, *p*=0.01; corrected for multiple comparisons), without a significant group × hemisphere interaction (F(40, 1)<0.01, *p*=0.98). Application of the same test to the measured activity within areas OPA, PPA, and RSC did not yield any significant effect of group and/or group × hemisphere interaction (**Table 5**). A separate ANOVA did not show any significant differences between the scene-selective activity measured in the strabismic vs. anisometropic individuals (*p>*0.45).

**Table 5.**
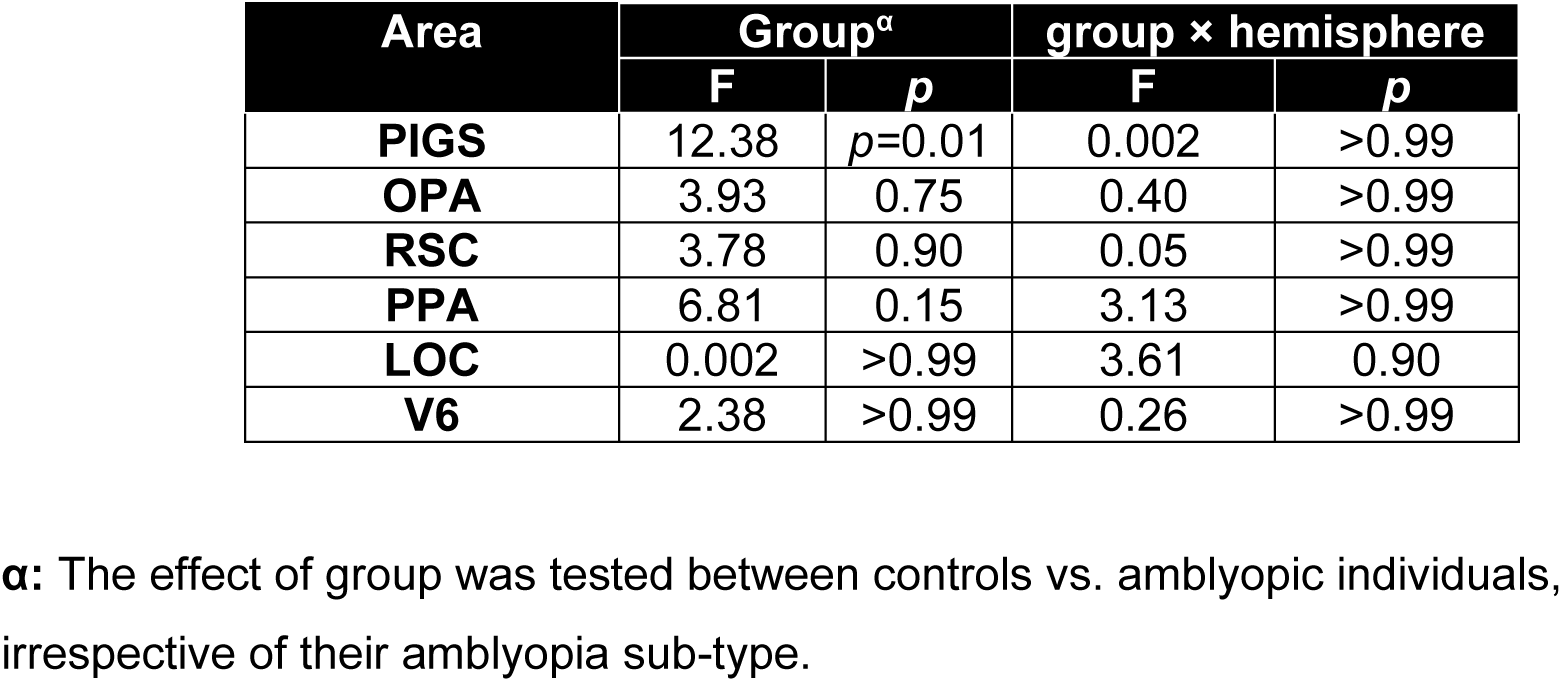
Between-group (amblyopic vs. control individuals) differences in the level of scene-selective activity.

As a control, we also measured the activity evoked within areas LOC, and V6 (see Methods). These control areas were selected based on their proximity to areas OPA and PIGS, respectively (Sulpizio et al., 2020; Kennedy et al., 2024). Tests in LOC and V6 showed no significant scene-selective activity difference between amblyopic and control individuals (**Table 5**). Thus, among those regions that showed scene-selective activity, reduced activity in amblyopia was mainly limited to area PIGS.

#### 3.3.5. Predictability of VFQ-39 scores based on the scene-selective responses

For the twenty individuals who participated in Experiments 1 and 2 (**Table *1***), we checked whether the reported scores for general vision, distance activities, and peripheral vision (based on VFQ-39) correlated with the level of scene-selective area activities based on the fMRI. Independent Pearson correlation tests showed a significant linear relationship between the reported score for general vision and the level of scene-selective activity within PIGS (R^2^=0.28, *p*=0.02), and OPA (R^2^=0.20, *p*=0.05) (Figure 7). General vision was not significantly correlated with activity in RSC or PPA (*p*>0.08; R^2^<0.15). Importantly, we did not find any significant correlations between measures of interocular visual acuity difference, binocular visual acuity, and stereoacuity with scene-selective activity across different ROIs (*p*>0.20; R^2^<0.10). These results suggest that the correlation between the scene-selective response and perceived visual function reflects the higher-order visual functional deficits of amblyopia rather than those of discrete spatial or interocular disparity resolution.

**Figure 7.**
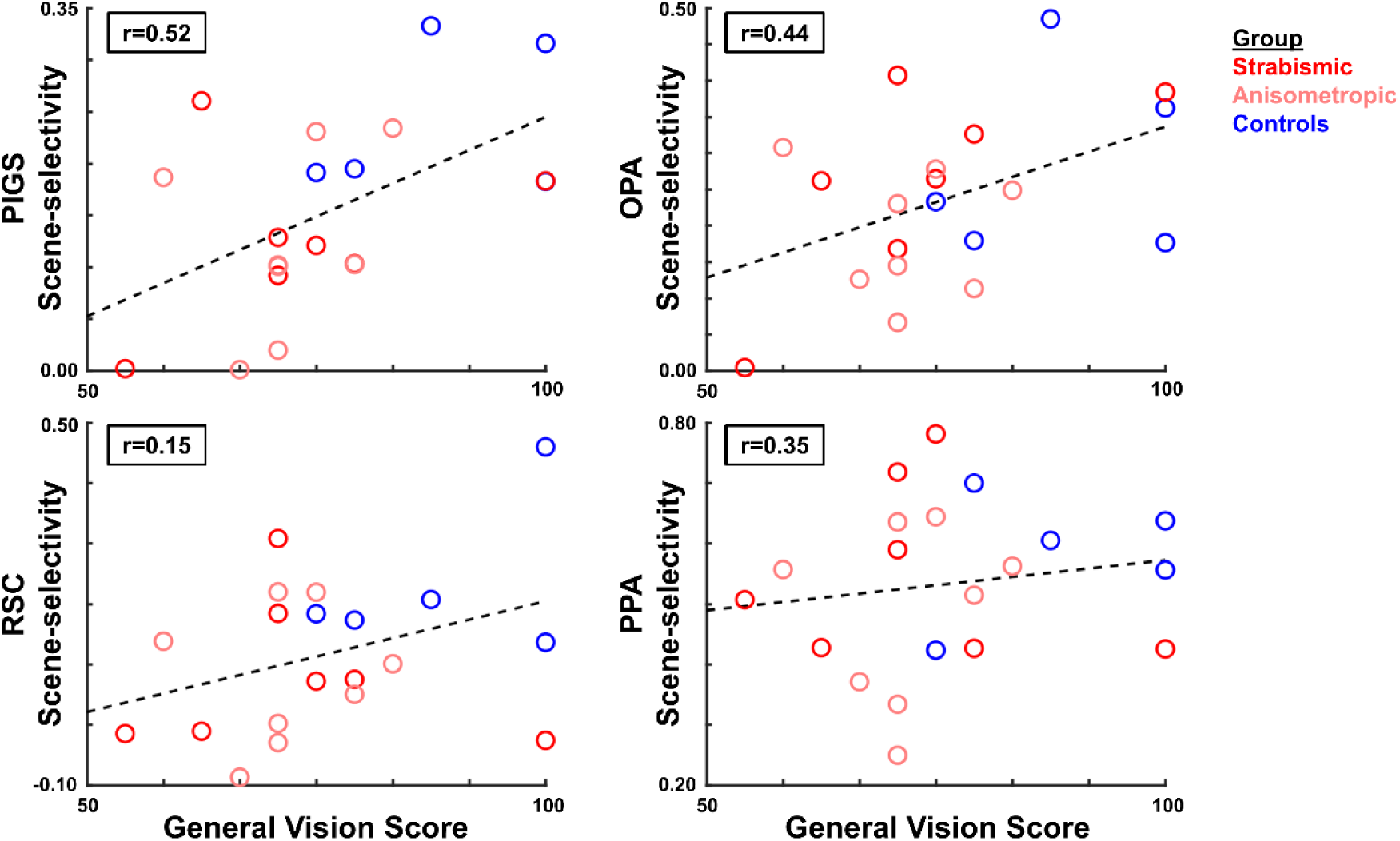
Correlation between the level of scene-selective activity and the reported score for general vision (VFQ-39 subscale). Across the scene-selective areas, we found a significant correlation only within areas PIGS and OPA. In each panel, each point represents data from one individual participant, averaged over the two hemispheres.

### 3.4. Experiment 3 – Object-selective cortical responses

Experiment 3 tested whether the amblyopic subjects also show a decreased object-selective activity, and if so, whether this reduction contributed to the observed decrease in the level of scene-selective activity in area PIGS. Accordingly, eighteen amblyopic individuals plus seventeen controls (selected from those who participated Experiment 2 based on their willingness to continue the scans) were scanned to measure their brain activity in response to intact vs. scrambled objects (see Methods and **Table *1***).

#### 3.4.1. Head position stability

As in Experiment 2, we compared the level of head motion during scanning, between control vs. amblyopic participants. A t-test applied to the measured level of head motion did not yield a significant difference between the two groups (t(33)=0.27, *p*=0.80), suggesting that head position was comparably stable between the two groups. Nevertheless, as in Experiment 2, we included this nuisance co-variate in all analyses, to eliminate residual effects of head motion.

#### 3.4.2. Group-averaged object-selective response

In both groups, object-selective activity was detected within a large portion of the extra-striate visual cortex, including the scene-selective areas PIGS, OPA, RSC, and PPA, plus object-selective area LOC (Figure 8A-B). In contrast to scene-selective activity maps, here, the vertex-wise between-group comparison did not show any significant difference between amblyopic and control participants across the scene-selective areas (Figure 8C-D) and also within area LOC. Thus, object-selective responses were comparable between the two groups.

**Figure 8.**
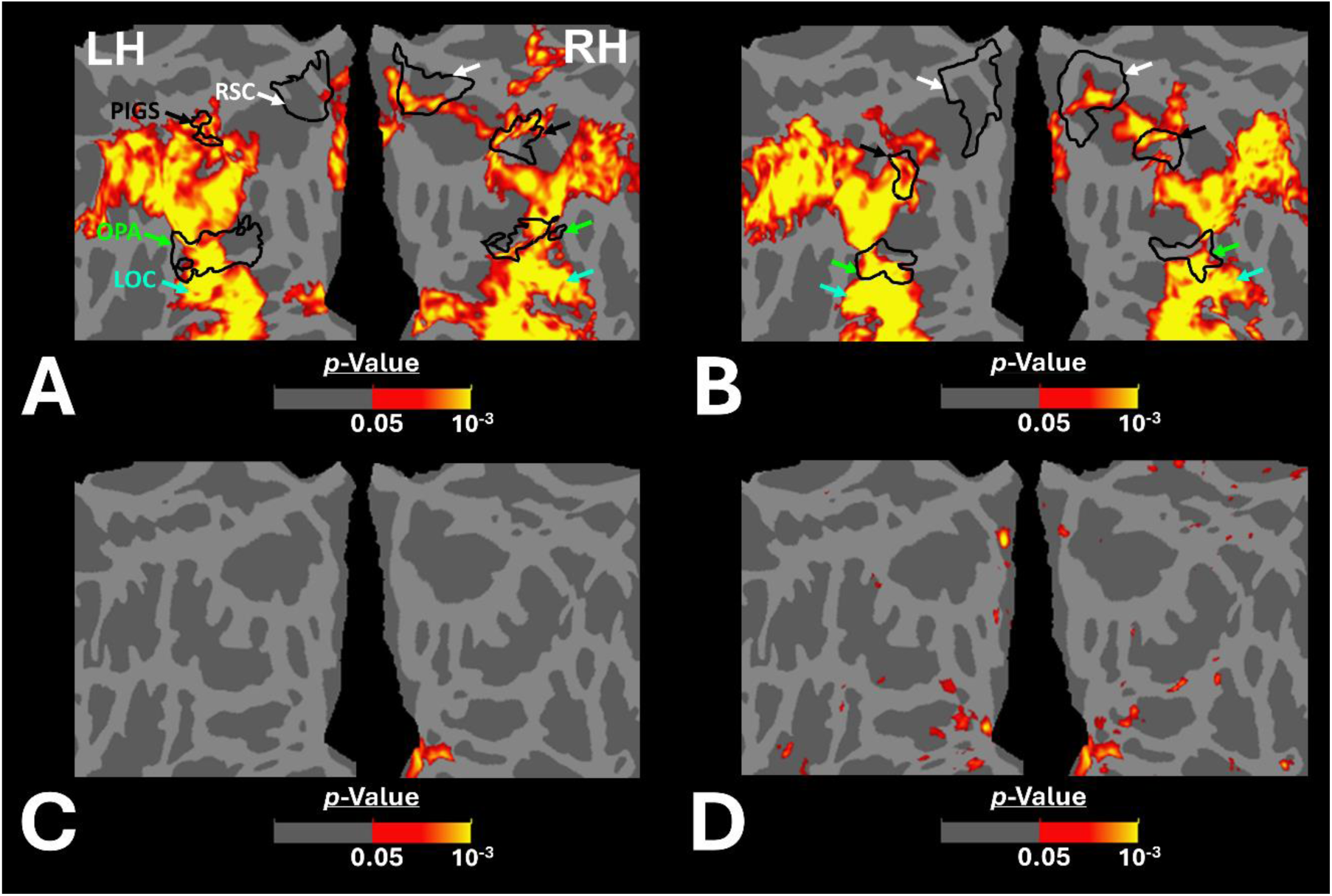
The group-averaged object-selective activity (intact object > scrambled object) in amblyopic and control participants across the occipito-parietal region. Panels A and B show the activity maps in controls and amblyopic individuals, respectively. In contrast to the scene-selective activity map within the same region (Figure 2) the overall pattern of object-selective activity appears to be comparable between the two groups, even within the posterior intraparietal region. Panels C and D show the between-group object-selective activity differences, with and without correction for multiple comparisons, respectively. Here again, we did not find any significant difference between the two groups.

#### 3.4.3. The amplitude of the object-selective activity across the amblyopic and control subject

Within the scene selective areas, we further compared the object-selective response, between amblyopic and control participants, using the more sensitive ROI-based analysis (Figure 9). Consistent with the group-averaged activity maps, results of this test did not yield any significant effect of group (*p*>0.72; corrected for multiple comparisons) and/or group × hemisphere interaction (*p*>0.48). These results suggest that the impact of amblyopia on the activity within the scene-selective areas was limited to the evoked response to scenes, and this effect was not attributable to altered object processing.

**Figure 9.**
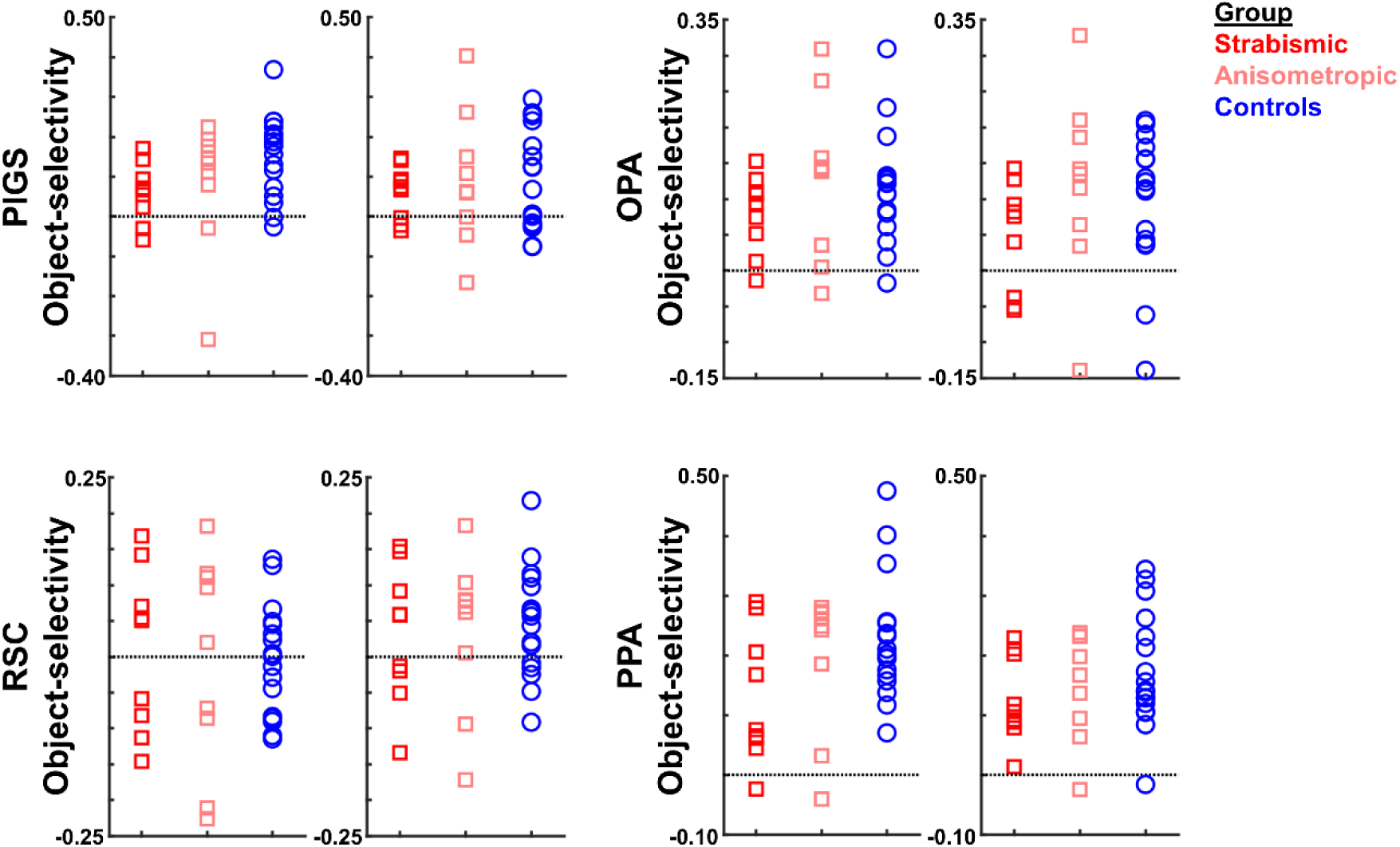
The level of object-selective activity (intact objects – scrambled objects) was measured across areas scene-selective areas PIGS, OPA, RSC, and PPA. Consistent with the activity maps, none of these areas showed any significant object-selective activity difference between amblyopic individuals and controls. All details are similar to Figure 6.

## 4. Discussion

The results of this study directly suggest that amblyopia impacts scene processing within PIGS, a scene-selective area located within the posterior intraparietal gyrus (Kennedy et al., 2024). We found that scenes (but not single objects) evoked weaker selective activity in PIGS (but not OPA, RSC and PPA) in amblyopic participants compared to controls. Correlation of scene-selective activity in this area with the reported score for general vision – a subscale in VFQ-39 questionnaire for quality of life – suggests meaningful functional relevance of this finding.

### 4.1. Amblyopia impacts on higher-order visual processing

Originally, amblyopia studies were mainly focused on area V1 because neurons within this cortical area show strong ocular preference (Hubel and Wiesel, 1962). According to early studies, amblyopia (especially in more severe forms) was associated with a decrease in the number of neurons that respond preferentially to the amblyopic eye (Crawford and Von Noorden, 1979; Crawford et al., 1996; Smith III et al., 1997b; Kiorpes et al., 1998). Later studies showed that amblyopia is also associated with a decrease in the number of disparity-selective neurons that are distributed over the extrastriate visual areas such as V2 (Nakatsuka et al., 2007; Bi et al., 2011). Still, to the best of our knowledge, no previous studies had tested the impact of amblyopia on higher-level visual areas involved in more complex visual processing such as object encoding.

Our current results suggest that the impact of amblyopia on the development of the visual cortex extends well into the higher-level visual areas that respond selectively to scenes. For one thing, stimuli were presented binocularly rather than monocularly to minimize the potential impacts of visual acuity difference between amblyopic individuals and controls. Moreover, the stimuli did not contain any stereo cues and did not induce any coherent motion – two visual features whose encoding is impaired in amblyopic participants (McKee et al., 2003; Aaen-Stockdale et al., 2007; Aaen-Stockdale and Hess, 2008; Levi et al., 2015). Nevertheless, deficient PIGS activity was evoked by scenes, but not non-scene objects. Together, our findings indicate that amblyopia selectively impacts scene processing within this region.

### 4.2. Why PIGS but not the other scene-selective areas?

PIGS is located within the posterior intraparietal gyrus, an area populated with motion- and stereo-selective sites (Tootell et al., 2022; Kennedy et al., 2023). It also contributes to ego-motion encoding within naturalistic environments, a perceptual process that relies heavily on depth, motion coherency (optic flow), and ego distance estimation. Previous psychophysical studies have shown that all these visual functions (i.e. depth, motion coherency and ego distance perception) are, at least to some extent, impaired in amblyopic individuals (McKee et al., 2003; Melmoth and Grant, 2006; Aaen-Stockdale et al., 2007; Aaen-Stockdale and Hess, 2008; Carlton and Kaltenthaler, 2011; Grant and Moseley, 2011; Levi et al., 2015; Ooi and He, 2015). Thus, in the absence of normal visual input, experience-dependent development of PIGS could be disrupted in amblyopia. Our findings of decreased scene-selective activity within the PIGS in amblyopic individuals supports a neurodevelopmental component to abnormal amblyopic scene processing. Such a developmental disorder may be even worsen over time in the absence of engagement in visually guided tasks that rely on ego-motion (Sá et al., 2021; Harrington et al., 2023), which also reduces the level of feedback to these regions.

In contrast to PIGS, PPA and RSC do not respond to ego-motion (Hacialihafiz and Bartels, 2015; Kamps et al., 2016; Jones et al., 2023; Kennedy et al., 2024). Rather, PPA and RSC are involved in scene recognition (Epstein et al., 2007; Park and Chun, 2009) and layout representation (Wolbers et al., 2011), respectively. Area OPA also appears to respond to simpler forms of motion, whose encoding is less affected by amblyopia. However, considering the functional connection between scene-selective areas (Baldassano et al., 2013; Nasr et al., 2013; Baldassano et al., 2016), it may be possible to see more extended between-group differences during complex, scene-related tasks that rely simultaneously on multiple visual cues.

### 4.3. Weaker input from the amblyopic eye is not the sole cause of scene-selective activity decrease in PIGS

By relying on their fellow eye, amblyopic participants usually show comparable binocular visual acuity relative to normally sighted individuals. However, it can still be argued that stimulation of the amblyopic eye likely evokes a weaker visual response, compared to either their fellow eye or the non-dominant eye in controls (Conner et al., 2007; Dorr et al., 2019; Nasr et al., 2024). Considering this, one may suggest that the activity decrease in PIGS is due to a weaker bottom-up input from the earlier visual areas to PIGS.

Two key results from our study argue against this explanation. First, intact scene-selective activity in other areas, including OPA, PPA, RSC, and V6, suggests that the decreased visual input may not be the sole reason for this phenomenon. Notably, the impact of decreased visual input from the amblyopic eye is expected to be stronger on OPA and V6 that contribute to earlier stages of scene processing compared to PIGS. Specifically, OPA overlaps with visual areas V3A/B and V7 (IPS0) and responds to simple visual cues such translational motion (Nasr et al., 2011; Silson et al., 2016), whereas PIGS is located adjacent to areas IPS2-4, and only responds to more complex visual cues such as optic flow caused by ego-motion in naturalistic scenes but not random dots (Kennedy et al., 2024). Area V6 is located adjacent to PIGS and responds selectively to scenes (Pitzalis et al., 2015; Sulpizio et al., 2020; Kennedy et al., 2024). But in contrast to PIGS, it responds strongly to optic flow caused by moving dots (Pitzalis et al., 2010; Kennedy et al., 2024). Moreover, the differential response to the stimulation of amblyopic vs. fellow eye is stronger in lower-rather than the higher-level visual areas (Nasr et al., 2024). Thus, the impact of this phenomenon is expected to be stronger in OPA and V6 compared to PIGS. However, contrary to this expectation, our results showed that the between-group scene-selective activity difference was mostly limited to PIGS. Second, we found that non-scene, object-evoked activity in all examined regions (including PIGS) was not impacted by amblyopia, as would be expected if reduced visual gain drove differences in these higher-order regions. Thus, decreased visual input from lower-level visual areas cannot explain the weaker scene-selective response in the PIGS.

### 4.4. Potential contribution of attentional deficits in amblyopia

Degraded visual attention in amblyopia has been previously reported by others (Ho et al., 2006; Hou et al., 2016; Verghese et al., 2019). In this study, we reduced the influence of attention on the level of scene-selective responses by instructing the participants to perform an orthogonal task (i.e., detection of color changes in the fixation spot). Nevertheless, it might be argued that uncontrolled attentional demand contributed to between-group activity difference that we found in PIGS. However if this were the case, we would have expected to see the same effect in the other scene selective areas, such as PPA - in which attention to scenes increases the level of scene-selective activity (O’craven et al., 1999; Nasr and Tootell, 2012; Baldauf and Desimone, 2014). In the absence of such an effect, the potential impacts of amblyopia on attention are unlikely to significantly contribute to the decreased scene-selective activity in PIGS.

### 4.5. Monocular vs. binocular visual stimulation

At least for individuals with intact binocular visual acuity, scene perception is a binocular task. Although it has been shown that scene perception impairments in amblyopic individuals are equally detectable under monocular and binocular conditions (Mirabella et al., 2011), future studies could test whether scene presentation to the amblyopic vs. non-amblyopic eye may evoke a differential response in PIGS. However, the relevance of such a hypothetical study is arguable, since everyday scenes are typically viewed binocularly, whenever possible.

### 4.6. Limitations

Amblyopia influences many aspects of visual perception, from stereopsis to motion coherency. To distinguish the impact of amblyopia on scene perception and the underlying neuronal processing from its impact on depth and motion coherency encoding, we designed a paradigm based on using 2D stimuli that did not induce any coherent motion. While this paradigm serves to isolate the impact of amblyopia on scene-selective processing, our approach may underestimate the impact of amblyopia on scene perception as experienced by amblyopic individuals in their daily lives and weakens the correlation between the level of evoked brain activity and self-reported visual functional scores. However, the correlation between the scene-selective activity and general vision score relationships in PIGS and OPA Figure 7 argues that the relationship is maintained across the functional spectrum in our sample.

Natural scene perception relies on input from the peripheral visual field (Levy et al., 2001; Hasson et al., 2002; Levy et al., 2004; Larson and Loschky, 2009). Thus, our restriction of visual stimuli to the central 20 degrees of the visual field may miss important contributions of more peripheral scene cues in a more immersive environment. On the other hand, central stimulation mitigates potential confounding by differences in binocular visual field sensitivity (perhaps more often seen in strabismus).

Our experimental design did not instruct the participants to explicitly categorize the stimuli into scenes vs. non-scenes, and/or to discriminate scene stimuli from each other. These constrains limited our analysis to activity evoked within the sensory visual areas, even though we reported multiple cortical sites in association brain area, including TPJ and DLPFC, in which we found between-group differences in the level of scene-selective response. The existence of these more anterior cortical sites suggests that amblyopia impacts may extend well beyond the sensory regions into the association cortical areas that control different aspects of the human behavior, from attention control (Corbetta et al., 2000; Shulman et al., 2007) to perceptual decision-making (Heekeren et al., 2004; Philiastides et al., 2011). Such an extension may become even more apparent when participants are involved in an active scene-related task (e.g. navigation).

## Conclusion

Our results show that the impact of amblyopia extends beyond the early retinotopic visual areas into cortical regions involved in scene processing. The results also highlight the likelihood that amblyopia affects the function of association brain regions such as LPMA, TPJ and DLPFC. Future studies employing more realistic, immersive stimuli (and tasks) that better resemble daily visual experiences could more comprehensively highlight the functionally relevant neural consequences of amblyopia in higher order visual areas.

## Acknowledgments

This work was supported by NIH NEI (grants R01 EY017081 and R01 EY030434), and by the MGH/HST Athinoula A. Martinos Center for Biomedical Imaging. Crucial resources were made available by a NIH Shared Instrumentation Grant S10-RR019371. We thank Dr. Adam Steel and Dr. Caroline Robertson for sharing data and insightful discussions.

